# Essential oil terpenes may inhibit steroidogenic cytochrome P450 activities

**DOI:** 10.1101/2023.10.31.564977

**Authors:** Katyayani Sharma, Angelo Lanzilotto, Jibira Yakubu, Søren Therkelsen, Clarissa Daniela Vöegel, Therina Du Toit, Flemming Steen Jørgensen, Amit V. Pandey

## Abstract

Endocrine-disrupting chemicals (EDCs) may impact the development of Prostate Cancer (PCa) by altering the steroid metabolism. Although their exact mechanism of action in controlling tumor growth is not known, EDCs may inhibit steroidogenic enzymes such as Cytochrome P450 c17 (CYP17A1) or aromatase (CYP19A1) involved in the production of Androgens or Estrogens. High levels of circulating androgens are linked to PCa in men and Polycystic Ovary Syndrome (PCOS) in women. Essential Oils or their metabolites (EOs) like lavender oil and tea tree oil have been reported to act as potential EDCs and contribute towards sex steroid imbalance in case of prepubertal gynecomastia in boys and premature thelarche in girls due to the regular exposure to lavender-based fragrances among Hispanic population. We screened a range of EO components to determine their effects on CYP17A1 and CYP19A1 Computational docking was performed to predict the binding of EOs with CYP17A1 and CYP19A1 and functional assays were done using the radiolabeled substrates or Liquid Chromatography high-resolution Mass Spectrometry and cell viability assays were carried out in LNCaP cells. Many of the tested compounds bind close to the active site of CYP17A1, and (+)-Cedrol had the best binding with CYP17A1 and CYP19A1. Eucalyptol, Dihydro-β-Ionone & (-)-α-pinene showed 20% to 40% inhibition of dehydroepiandrosterone production; and some compounds also effected CYP19A1. Extensive use of these EOs in various beauty and hygiene products is common, but only a limited knowledge about their potential detrimental side effects exists. Our results suggest that prolonged exposure to some of these essential oils may result in steroid imbalances. On the other hand, due to their effect on lowering androgen output, ability to bind at the active site of steroidogenic cytochrome P450s, these compounds may provide design ideas for the novel compounds against hyperandrogenic disorders such as PCa and PCOS.

## 1. Introduction

Essential Oils (EOs) are a complex mixture of volatile compounds extracted from aromatic plant tissues with a characteristic “essence” or smell [1]. Pure extracts of EOs are obtained through different methods such as steam distillation, solvent extraction, and hydro distillation [2]. The chemical composition of EOs can vary depending on the origin and species of the plant, climate, and extraction method [3]. Two major constituents of EOs are terpenes and terpenoids [4]; some examples of terpenes found in EOs are cineol (eucalyptol), linalool, pinene, limonene, thujene, bisabolene, caryophyllene, p-cymene, camphor, neral, menthol, and geraniolwhile aromatic compounds consist of carvacrol, thymol, cinnamaldehyde, eugenol, and estragole [5]. Owing to their charestric fragrance, EOs are extensively used in many cosmetics and hygiene products [6] [7]. Due to their anti-microbial, antibiotic, antiviral, antioxidant, and anti-inflammatory properties, they have been part of traditional therapies and herbal medicines [8] [9] [10] **(Figure 1)**. Being “natural” in origin, EOs are often considered as safer substitutes of chemical drugs that may have adverse side effects [11]. However, in addition to their therapeutic role, EOs might function as potential Endocrine Disrupting Chemicals (EDCs).

**Figure 1:**
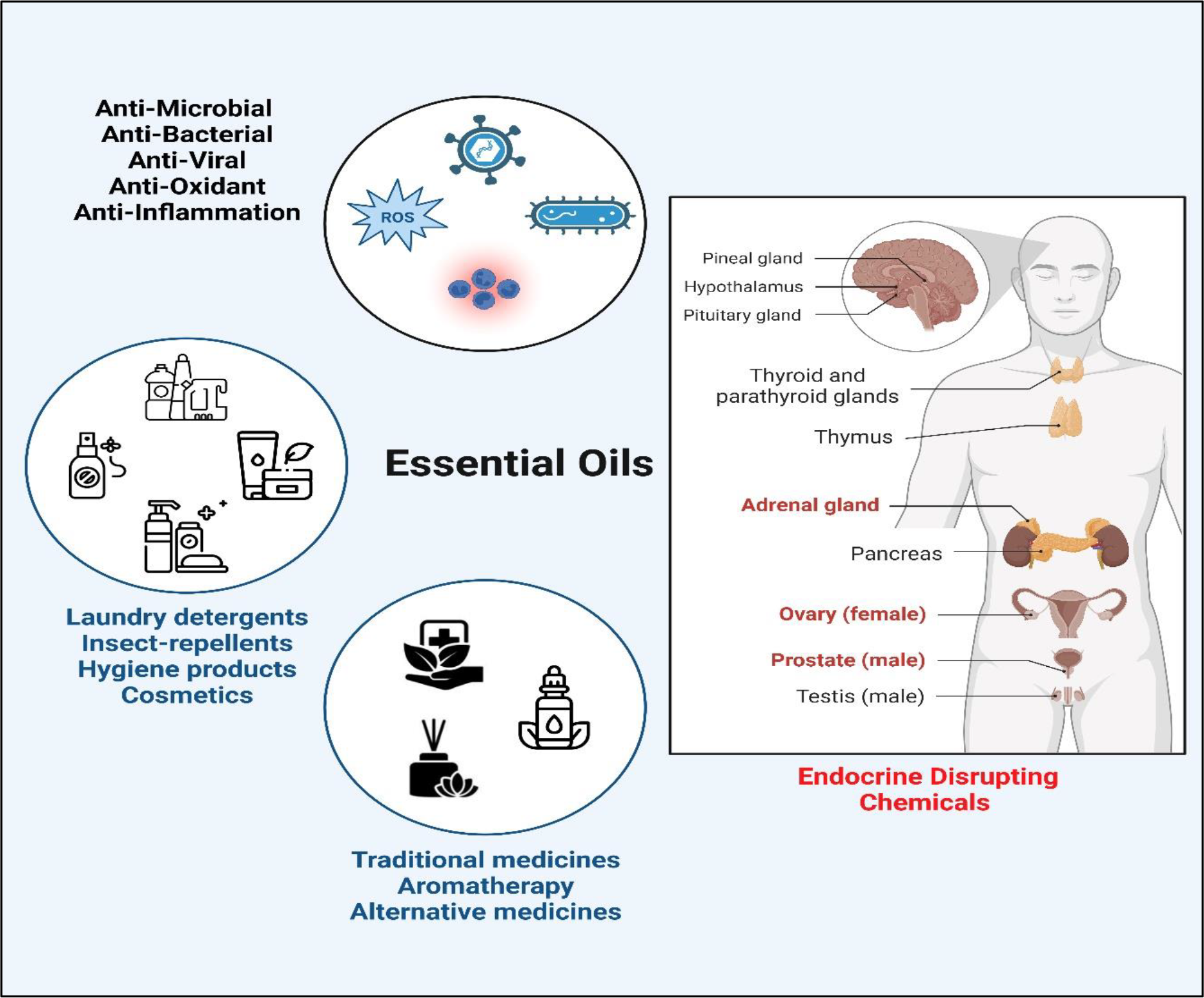
An overview of biological activities of essential oil components.

EDCs are chemical substances that can alter endocrine function by interfering with steroid metabolism resulting in hormonal imbalance in the body. Abnormal levels of steroids, especially sex steroids, can cause detrimental effects on sexual development and possess an increased risk of infertility [12], though the exact mechanism of action of EDCs is not fully known. Clinical case reports have linked prepubertal gynecomastia in boys and premature thelarche in girls to prolonged use of lavender and tea tree oil-based fragrant products which resolved upon cessation of the products. Moreover, studies in human breast cancer cell lines showed estrogenic and anti-androgenic activity of some EOs [8] [13].

In humans, androgens are primarily produced in the male testis, female ovaries, and adrenal glands (**Figure 2**). Androgens control male sexual traits and development as well as influence female sexual behavior. The zona reticularis of the adrenal cortex produces dehydroepiandrosterone (DHEA) and its sulfate DHEA(S). DHEA acts as a precursor to produce androgens (testosterone and androstenedione). The first rate-limiting step in the biosynthesis of all steroid hormones is the cleavage of the cholesterol side chain by the mitochondrial P450 enzyme CYP11A1 system (CYP11A1-FDX-FDXR) also called P450scc, to convert cholesterol into pregnenolone (Preg) [14]. Further in multi-enzymatic steps, Preg is converted into mineralocorticoids, glucocorticoids, and androgens.

**Figure 2:**
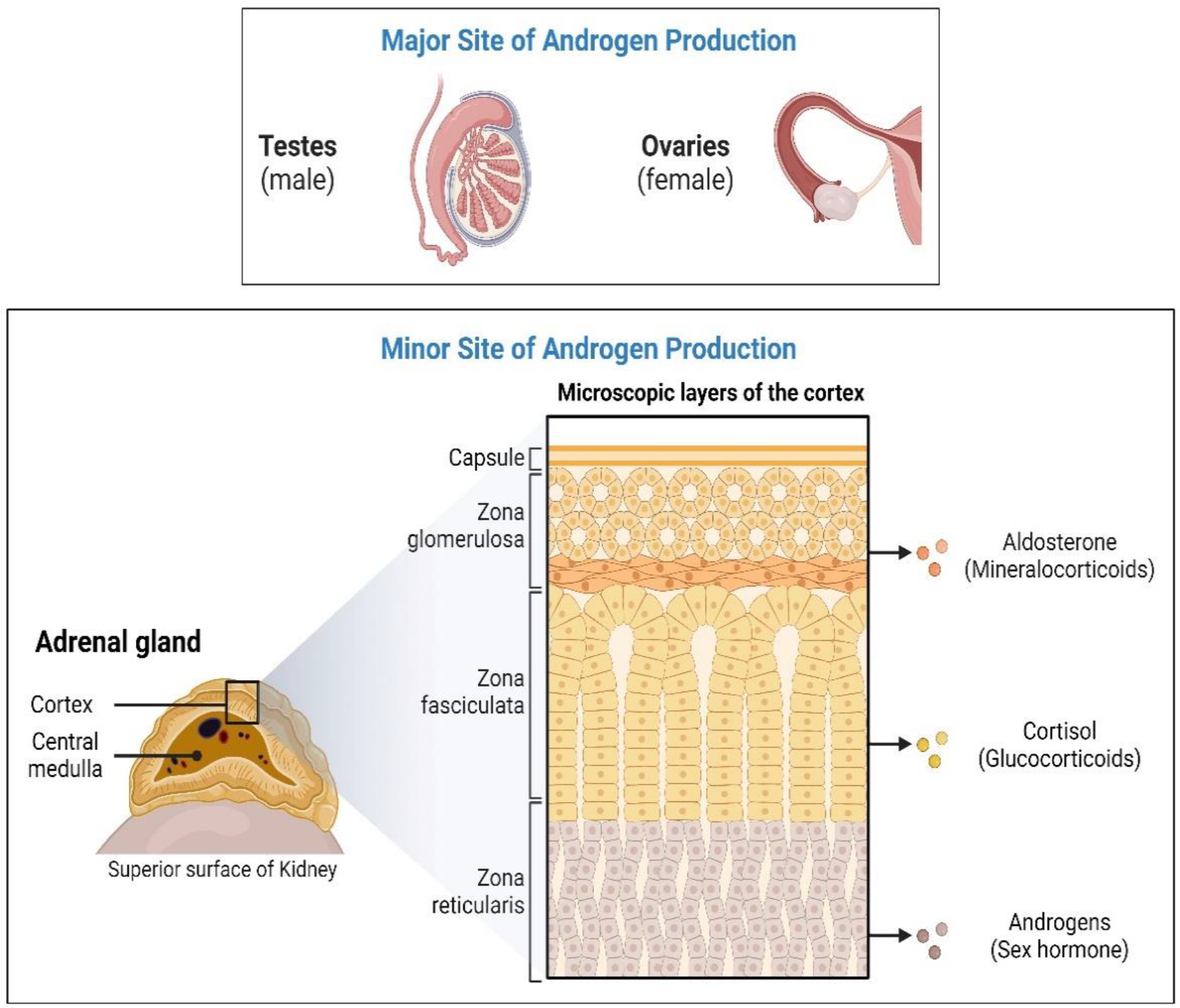
Androgen production in humans. In humans, androgens are mainly produced in the adrenal glands, male testis and female ovaries. Sexual traits and development as well as influence on female sexual behavior may be regulated by androgens. Androgen precursors, dehydroepiandrosterone (DHEA) and its sulfate DHEA(S) are produced in the zona reticularis of the adrenal cortex which are then converted into active androgens by the action of a series of steroid metabolizing enzymes.

The enzyme P450c17 (CYP17A1, Cytochrome P450 17alpha-monooxygenase), encoded by *CYP17A1* gene [15, 16] is an essential enzyme that plays a vital role in adrenal androgen production [16, 17]. The CYP17A1 localized in the endoplasmic reticulum can catalyze both 17α-hydroxylase and 17,20 lyase reactions [18]. This characteristic dual activity is conferred through post-translational regulation of CYP17A1 protein. Especially, the 17, 20 lyase activity of CYP17A1 is supported by at least three factors. First, the amount of P450 Oxidoreductase (POR) for electron transfer [19, 20], second, the presence of allosteric activator microsomal Cytochrome B5 (CYB5) [21, 22], and third, the phosphorylation of the CYP17A1 protein at serine/threonine residues [23] [22, 24] [25] [26]. Understanding the regulatory mechanisms of 17,20 lyase activity is important for the understanding of hyperandrogenic disorders such as premature, exaggerated adrenarche, PCa, and PCOS [27, 28].

Epidemiological studies suggest that EDCs may function as hormone mimics and bind to nuclear receptors to elicit altered expression of genes involved in the development and progression of Prostate Cancer (PCa) [29] (**Figure 3**). Prostate tumor cells are driven by Androgens binding to the intracellular Androgen Receptor (AR). The AR acts as a transcriptional activator for the expression of genes responsible for the growth and survival of the tumor [30]. These studies suggest that the possible role of EOs could be either in direct inhibition or co-activation of certain steroidogenic enzymes or in the regulation of gene expression of these enzymes resulting in abnormal androgen levels in the body. High levels of circulating androgens are linked to both PCa and Polycystic Ovary Syndrome (PCOS) [31]. The anti-androgenic property of EOs could be exploited to target androgen production in the treatment of PCa and PCOS.

**Figure 3:**
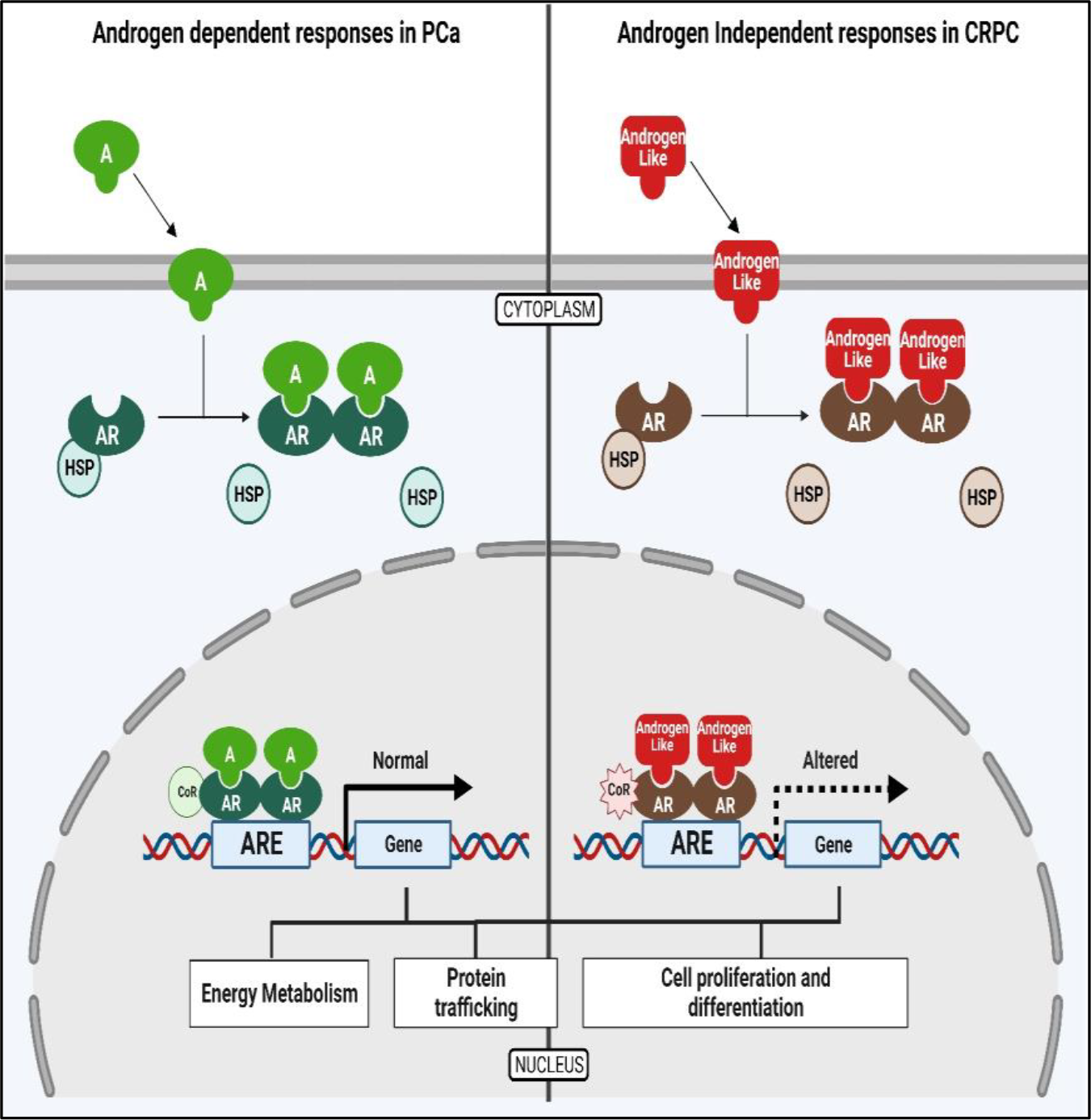
Prostate cancer cells are driven by the binding Androgens to the Androgen Receptor (AR). The AR then acts as a transcriptional activator for the expression of genes that are responsible for the growth and progression of the prostate cancer cells.

Considering the potential role of essential oils as EDCs, we explored the potential of several common essential oil components on human steroid metabolizing enzymes CYP17A1 and CYP19A1 for potential effects on androgen and estrogen production in humans and as potential structural leads for design of novel chemicals targeting these enzymes in hormone dependent cancers. We docked 53 terpene compounds that are naturally present in essential oils, into the structures of steroid-metabolizing enzymes CYP17A1 and CYP19A1 in order to estimate binding affinity and possible binding modes and sites to identify possible endocrine disrupting properties of these compounds.

## 2. Materials and Methods

### Terpenes

The terpenes used in the experiments cannot be called essential oils, as they do not exist as mixtures of compounds but are commercially available as single chemical entities (sometimes multiple isomers coexist because the separation process can bring purity only to a certain point). The terpenes we sourced are of mixed natural/synthetic origin depending on which provided higher purity. Their isolation usually consisted of an essential oil first collected through steam distillation or alcoholic extraction of the dry plant/flower mass, and then fractional distillation to collect the main components.

All terpenes were sourced from Sigma Aldrich, unless otherwise indicated, and the (individual product purity was between 90-99%) product codes were: (-)-α-Pinene (305715, (purity 99%), (+)-α-Pinene (268070, purity 99%), α-Ionone (I12409, purity 90%), Benzaldehyde (B1334, purity 99%), p-Anisaldehyde (A88107, purity 98%), 1,4-Cineole (W365820, purity 95%), Isoamyl acetate (W205532, purity 97%), Octyl acetate (W280607, purity 98%), Benzyl acetate (B15805, purity 99%), Propyl acetate (133108, purity 99%), β-Pinene (402753, purity 99%), Bisabolene (Alfa Aesar, A18724, mixture of isomers), (−)-α-Bisabolol (14462, purity 93%), 3-Carene (115576, purity 90%), (S)-(+) Carvone (435759, purity 96%), (+)-Cedrol (22135, purity 99%, sum of enantiomers), Cinnamyl alcohol (108197, purity 98%), p-Cymene (C121452, purity 99%), Dihydrocarvone (218286, purity 98%, mixture of isomers), Dihydro-β-ionone (W362603, purity 90%), Eucalyptol (C80601, purity 99%), Farnesol (W247804, purity 95%, mixture of isomers), Geraniol (163333, purity 98%), Methyl anthranilate (W268208, purity 98%), (R)-(+)-Limonene (183164, purity 98%), (+-)-Citronellal (27470, purity 95%), (R)-(-) Carvone (124931, purity 98%), (1R)-(-) Myrtenal (218243, purity 98%), Nerol (268909, purity 97%), Ocimene (W353977, purity 90%, mixture of isomers), (S)-(-)-Limonene (218367, purity 96%), Carvacrol (282197, purity 98%), (+)-Sabinene (W530597, purity 75%), (S)-(-) Perillyl alcohol (218391, purity 96%), Estragole (A29208, purity 98%), (-)-α-Terpineol (W304522, purity 96%), Terpinolene (W304603, purity 95%), Thymol (T0501, purity 98%), Vanillin (V1104, purity 99%), Methyl salicylate (M6752, purity 99%), α-Terpinyl acetate (W304799, purity 95%), α-Phelladrene (W285611, purity 85%), γ-Terpinene (223190, purity 97%).

### Molecular Docking Analysis

In a first attempt, 53 terpene compounds were docked into CYP17A1 and CYP19A1 using AutoDock VINA [32] [33]. Ligands co-crystallized with the PDB structures [34] in PDB IDs 4NKZ [35] and 3S79 [36-38] (CYP17A1 and CYP19A1, respectively) were removed, and the remaining protein structures were used for docking. Three-dimensional structures of the ligands were extracted from PubChem and prepared for docking using the LigPrep [39] function within Maestro [Schröinger Release 2022-3: Maestro, Schröinger, LLC, NY, 2021], removing possible salts and ensuring generation of possible ionization and tautomeric states at pH=7±1 using the Epik [40] setting. Prior to docking the compounds were subjected to a short energy minimization. As reference compounds, abiraterone and 17α-hydroxypregnenolone were docked into CYP17A1 as well and androstenedione into CYP19A1 and compared with known structures of CYP17A1 [35, 41, 42] and CYP19A1 [36-38, 43]. This yielded a global docking simulation including the whole protein structure. For each ligand, 25 docking runs were performed. The results were subjected to a cluster analysis with each cluster differing at least 5Å heavy atom RMSD, representing different possible sites and modes of binding.

In a further refinement of this process, the compounds were docked into not only CYP17A and CYP19A1, but also to CYP11A1 and CYP21A2 with GLIDE [v 5.8, Schröinger, LLC, NY, 2021] using both the SP and XP scoring functions [44, 45]. Subsequently, the best scoring poses for each compound for each enzyme and for each scoring function were extracted and analyzed and heat maps produced (cf. Figure 4).

**Figure 4:**
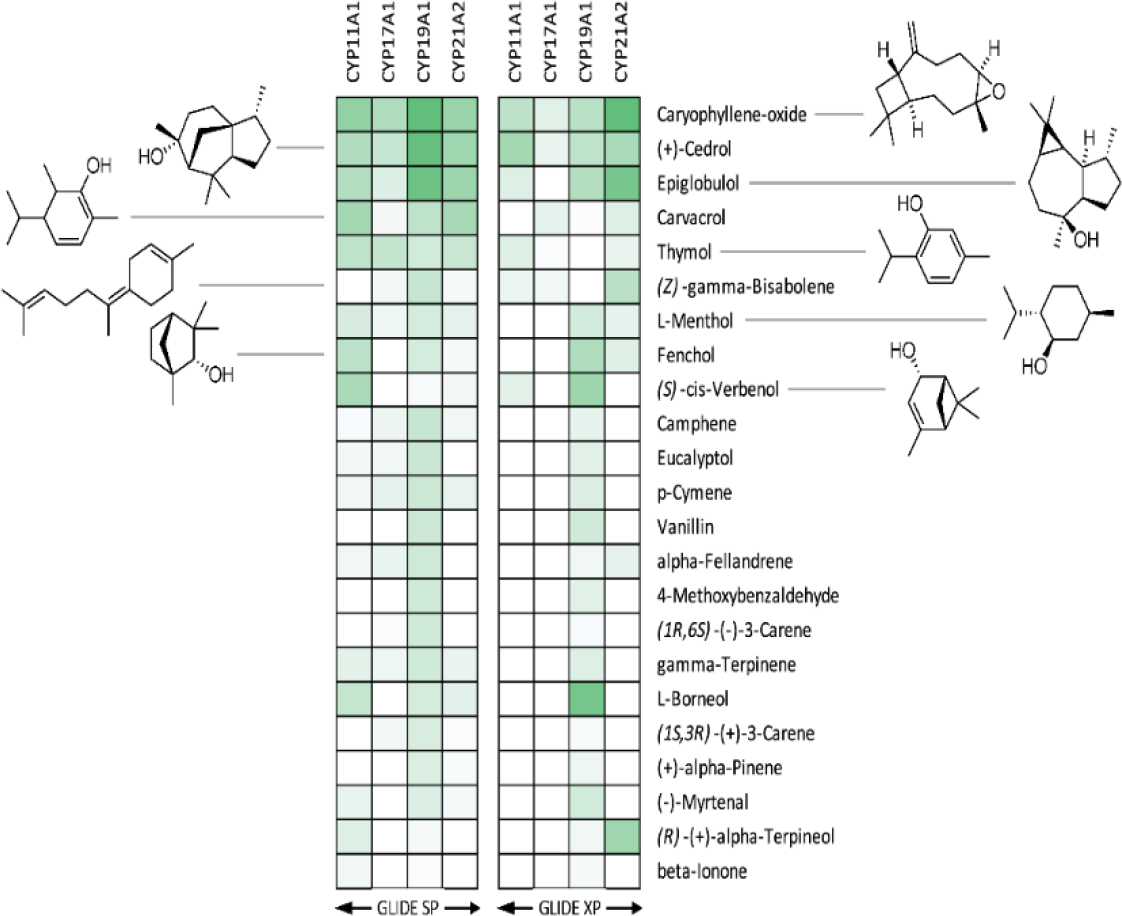
Heat maps of binding poses obtained after docking of terpenes from essential oils into steroid metabolizing CYP enzymes. Binding energies vary from -8.2 kcal/mol for the best binding poses (corresponding to dark green boxes) to -3.3 kcal/mol for the poorest binding poses (white boxes).

### Chemicals

Trilostane was obtained from the extraction of commercially available tablets as Modrenal® (Bioenvision, NY, USA). Abiraterone acetate was purchased from MedChemExpress®, Lucerna Chem AG (Lucerne, Switzerland). Commercially available drug, Anastrazole was purchased from AstraZeneca. Radiolabeled substrates, Progesterone [4-^14^C] (SA 55mCi/mmol; Conc. 0.1mCi/mL); 17α-Hydroxypregnenolone [21-^3^H] (SA 15Ci/mmol; Conc. 1mCi/mL) and Androstenedione [1β-^3^H(N)] (SA 24 Ci/mmol; Conc. 1mCi/mL) were obtained from American Radiolabeled Chemicals Inc. (St. Louis, MO, USA). Non-radiolabeled standard substrates, Pregnenolone; Progesterone; 17α-Hydroxypregnenolone; 3-(4,5-Dimethyl-2-thiazolyl)-2,5-diphenyl-2H-tetrazolium bromide (MTT); Resazurin sodium salt; Dimethyl sulfoxide (DMSO) and Dextran were purchased from Sigma-Aldrich^®^ (St. Louis, MO, USA). NADPH tetrasodium salt and Organic solvents such as Isooctane, Ethyl acetate, and Chloroform/Trichloromethane were acquired from Carl Roth® GmbH + Co. KG (Karlsruhe, Germany). Activated Charcoal was obtained from Merck AG (Darmstadt, Germany).

### Cell line and culture

The current standard model system to study molecular and biochemical mechanisms of steroidogenesis is the NCI H295R cell line [46] [47]. These cells express genes from all three zones of the adrenal cortex, providing an excellent system that closely reflects human adrenal physiology [14]. The human adrenocortical carcinoma cell line NCI H295R was obtained from the American Type Culture Collection (ATCC® CRL2128™), Manassas, VA, USA [46] [47]. Cells between passages 12-24 were cultivated in DMEM/Ham’s F-12 medium (1:1 Mix) supplemented with L-glutamine and 15 mM HEPES (Gibco™, Thermo Fisher Scientific, Waltham, MA, USA) along with 5% Nu-Serum I; 0.1% insulin, transferrin, selenium in form of ITS Premix (Corning™, Manassas, VA, USA); 1% Penicillin-Streptomycin (Gibco™, Thermo Fisher Scientific, Waltham, MA, USA) at 37°C in a humid atmosphere with a constant supply of 5% carbon dioxide to maintain the physiological pH. Human prostate cancer cell line, derived from metastatic site, left supraclavicular lymph node, LNCaP clone FGC (ATCC® CRL1740™) was cultured in RPMI-1640 Medium containing 2mM L-glutamine with 10mM HEPES, 1mM Sodium pyruvate, 10% Fetal Bovine Serum and 1% Penicillin-Streptomycin as supplements (Gibco™, Thermo Fisher Scientific, Waltham, MA, USA). For experiments, cells with passage numbers 12-30 were used as previously described [48].

### Cell Viability Assays

To determine the effect of test compounds on the cellular activity of human adrenal NCI H295R cells, MTT-based cell viability assay was performed [49] [50]. In a 96-well plate, about 30,000 cells per well were seeded with complete medium. After 24 hours, the medium was replaced with fresh medium and 10µM of test compounds were added. DMSO (less than 1% v/v) was used as vehicle control. 10µM Abiraterone was used as a positive control [51] [52]. 0.5mg/mL MTT reagent was added to the culture medium for another 4 hours. After the incubation, the medium was entirely replaced with DMSO to dissolve the formazan crystals. After 20 minutes, absorbance was measured at 570nm (SpectraMax M2, Bucher Biotec, Basel Switzerland). Percent viability is calculated with respect to the mean value of control samples.

For prostate cancer LNCaP cells, Resazurin-based Alamar blue assay was performed to evaluate the cell toxicity [49] [50]. Cells seeded at a cell density of 10,000 cells per well were treated with test compounds and the controls for 24 and 48 hours. After incubation, 0.05mg/mL Resazurin in Phosphate buffer was added. Cells were incubated for another 4 hours in dark at 37°C. Fluorescence was measured at an excitation wavelength of 550nm and an emission wavelength of 590nm. Percent viability is calculated with respect to the mean value of control samples (DMSO).

### CYP17A1 enzyme assays

The CYP17A1 enzyme assays were carried out according to well-established protocols [52] [53] in our laboratory. The NCI H295R cells were seeded overnight in a 12-well plate at a cell density of 0.5 X 10^6^ cells per well. Next day, 10µM of test compounds were added to respective wells containing fresh medium and incubated for 4 hours. Abiraterone and DMSO were used as reference and control respectively. To determine CYP17A1 hydroxylase activity, cells were treated with the [^14^C]-Progesterone at a concentration of 10,000cpm/1µM per well [22-24, 54]. Trilostane was added prior to the addition of test compounds and the substrate to block 3β-hydroxysteroid dehydrogenase activity [55]. Radiolabeled steroids were extracted from the media with help of Ethyl acetate and Isooctane (1:1 v/v) and separated through Thin Layer Chromatography (TLC) on a Silica gel coated aluminum plate (Supelco® Analytics, Sigma Aldrich Chemie GmbH, Germany) [56]. TLC spots were exposed to a phosphor screen and detected by autoradiography using Typhoon™ FLA-7000 PhosphorImager (GE Healthcare, Uppsala, Sweden). Radioactivity was quantified using ImageQuant™ TL analysis software (GE Healthcare Europe GmbH, Freiburg, Germany). Enzyme activity was calculated as a percentage of radioactivity incorporated into the product with respect to the total radioactivity.

Using similar treatment conditions, [21-^3^H]-17α-Hydroxypregnenolone (50,000cpm/1uM per well) was used as a substrate to analyze CYP17A1 Lyase activity. NCI H295R cells were treated with test compounds for 24hours before the addition of the substrate and Trilostane. Tritiated water release assay was performed [57] by measuring the conversion of 17OH-Preg into DHEA. Steroids in the media were precipitated using 5% activated charcoal/0.5% dextran solution. The enzyme activity was estimated with reference to the water-soluble tritiated by-product formed in an equimolar ratio with the corresponding DHEA. The radioactivity in the aqueous phase was measured by Liquid Scintillation counting (MicroBeta2® Plate Counter, PerkinElmer Inc. Waltham, MA, USA). The percent inhibition was calculated with respect to the control [58].

### Steroid Profiling

For steroid analysis, NCI H295R cells were treated in a similar way except that 1µM of the unlabeled substrate, Pregnenolone, was used instead of radiolabeled substrates for 4 hours. Steroids were measured by a liquid chromatography high-resolution mass spectrometry (LC-HRMS) method as previously described and validated [59]. Briefly, steroids were extracted from 500 µL cell media aliquots, plus 38 µL of a mixture of internal standards (at 3.8 nM each), using solid-phase extraction with an OasisPrime HLB 96-well plate. Samples were resuspended in 100 µL 33% methanol and 20 µL injected into the LC-HRMS instrument (Vanquish UHPLC coupled to a Q Exactive Orbitrap Plus, from Thermo Fisher Scientific) using an Acquity UPLC HSS T3 column (from Waters). Data from the mass spectrometer was processed using TraceFinder 4.0 (from Thermo Fisher). The lower limit of quantification (LOQ) for pregnenolone was 0.77 nmol/L, for DHEA it was 0.85 nmol/L, for DHEA-S it was 6.25 nmol/L and for 17OHP5 (quantified relative to the calibration of progesterone using a calculated response factor) it was 20 nmol/L.

### Aromatase (CYP19A1) Assay

Estrogens are synthesized from androgens through the action of the enzyme, Cytochrome P450c19a1 (CYP19A1, Aromatase) [60]. We used 40 µg of microsomal fraction from placental JEG-3 (Human Choriocarcinoma; ATCC^®^ HTB36^™^) cells in 100mM potassium phosphate buffer (pH 7.4) containing 100mM NaCl in a reaction mixture of 200 µL to carry out Aromatase enzyme activity assay. For determining the impact on aromatase activity, 10 µM of test compounds, DMSO as a negative Control, and Anastrozole (a known CYP19A1 inhibitor) as positive control were added to the reaction mixture. Tritium-labelled Androstenedione (∼30,000 cpm/µL/50nM) was used as the substrate to monitor the enzyme activity. The chemical reaction was initiated by the addition of reduced Nicotinamide adenine dinucleotide phosphate (NADPH) followed by incubation at 37°C with constant shaking for 1 hour. The reaction was stopped by the addition of Charcoal/Dextran solution. Enzyme activity was measured using a Tritiated water release assay as described earlier [53, 61].

### Statistical analysis

Calculations were done with Microsoft Excel and GraphPad Prism 3.0 (Graph Pad Software, Inc. San Diego, CA, USA). Data are represented as the mean of triplicate values from a single experiment or three independent sets of experiments. Dunnett’s multiple comparison ANOVA test was performed to determine the significant difference between the mean values of samples and the control. Error bars exhibit standard deviation from respective mean values. Significant p values were set as *p < 0.05 and **p < 0.01, ***p < 0.001.

## 3. Results

### Docking with CYP17A1 and CYP19A1

We performed computational docking and binding analysis of essential oil compounds against the three-dimensional crystal structures of multiple steroid metabolizing cytochrome P450 enzymes including CYP11A1, CYP17A1, CYP19A1 and CYP21A2.

Best binders were clustered into groups based on binding to multiple enzymes and subjected to detailed binding analysis. While we observed no significant selectivity (**Figure 4**), we identified the generally potential best binders for these CYP enzymes and made a preliminary look at the binding mode of the best poses. The conclusion is that a small number of compounds (Caryophyllene oxide, (+)-Cedrol and Epiglobulol), seems to bind reasonably well to the CYPs primarily by hydrophobic interactions. The compounds bind in the active sites of both CYP17A1 and CYP19A1 without coordinating directly to the Fe atom in the heme group (**Figure 5**). Docking studies revealed that (+)-Cedrol was among the best binding compounds to both CYP17A1 and CYP19A1 (Figure 6). Experimental results provided some context to the computational studies. Though it didn’t inhibit CYP17A1 activity, it caused 30% inhibition of CYP19A1 activity. Dihydro-β-Ionone showed 30% inhibition of both CYP17A1 and CYP19A1 activities. Eucalyptol and (-)-α-pinene showed 20% to 40% inhibition of CYP17A1. Although the inhibition was weaker than the reference compounds, it could be an inspiration for design of novel inhibitors, since the top scoring poses are rather globular compounds filling the cavity above the heme group in the CYPs.

**Figure 5:**
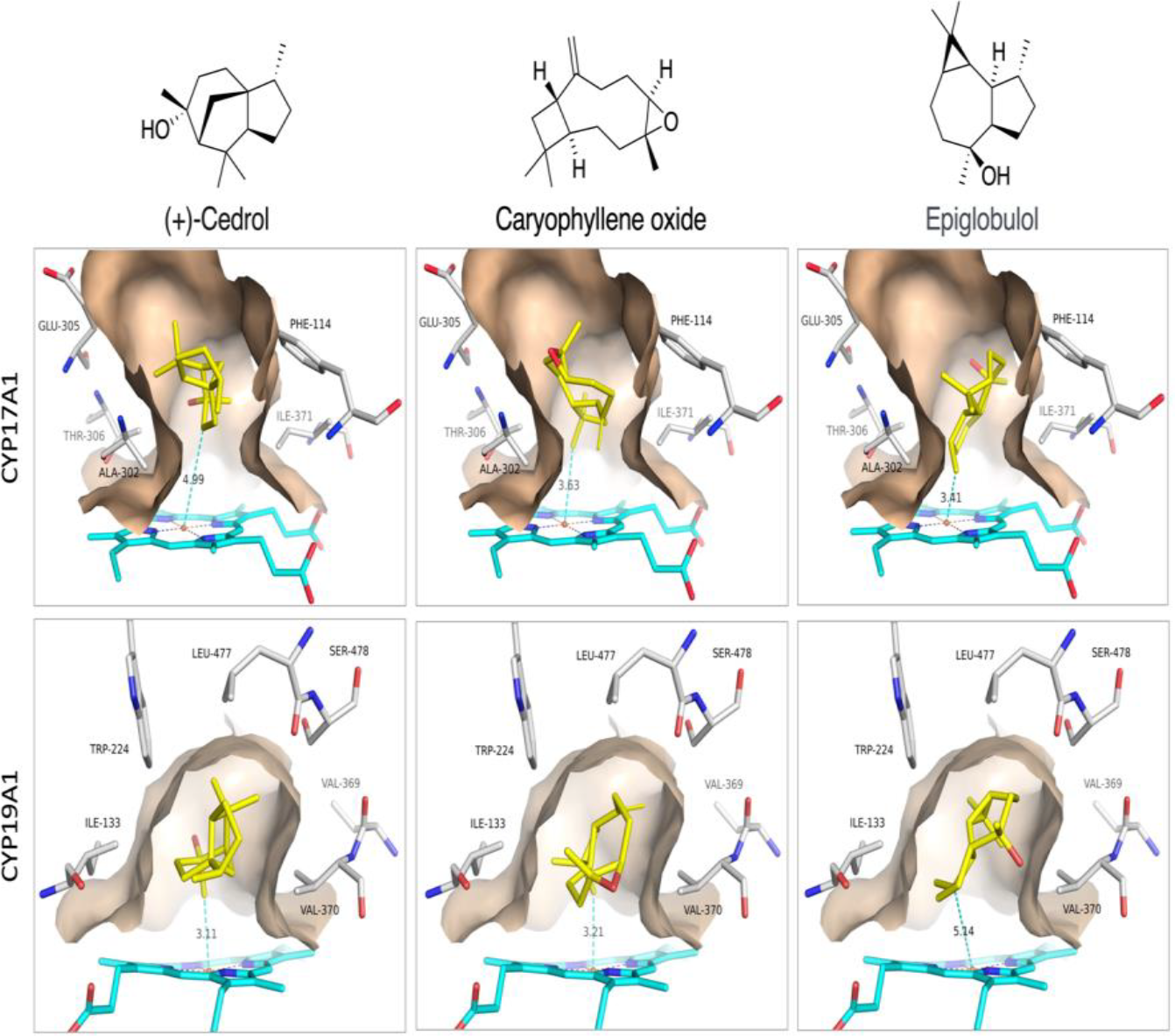
Proposed binding modes of (+)-Cedrol, Caryophyllene oxide and Epiglobulol to CYP17A1 and CYP19A1.

**Figure 6:**
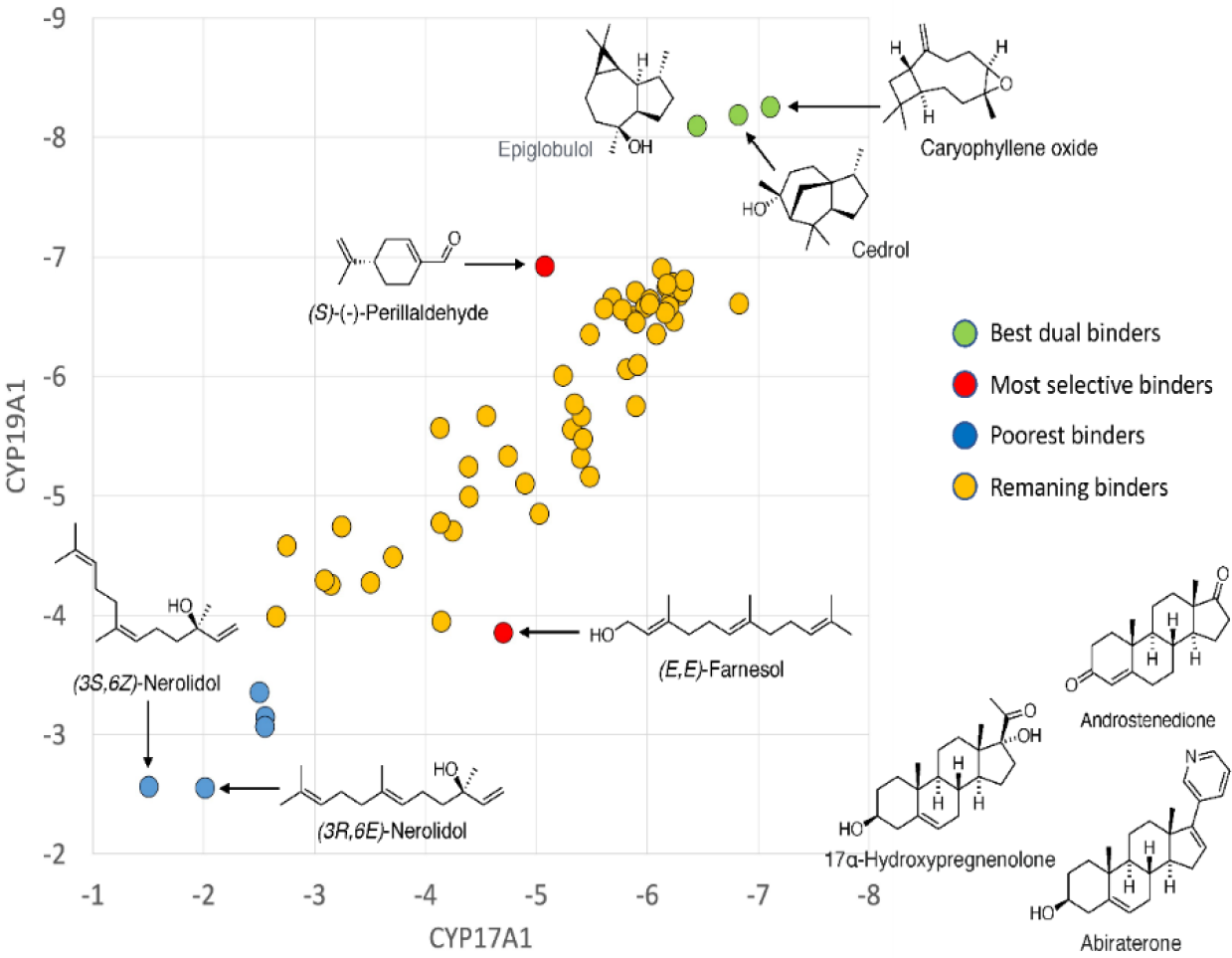
CYP17A1/CYP19A1 selectivity. Plot of binding energies to CYP17A1 and CYP19A1, respectively. Known substrates/inhibitors of CYP17A1 and CYP19A1 were used as controls.

### Effect on CYP17A1 Activity

Studies in human cell lines [8] [13] have shown that Lavender oil (LO) and Tea tree oil (TTO) act as hormone mimics for Estrogen receptors (ER) and antagonists for AR. Moreover, LO and TTO impacted the ER and AR-mediated regulation of several endogenous genes. Owing to these different mechanisms of action by LO and TTO, we screened several EO components including the ones found in TTO and LO. In the initial screening of 50 test compounds against CYP17A1 hydroxylase activity, we found no significant effect in NCI H295R cells treated with 10 µM of compounds for 4 hours (**Figure 7**).

**Figure 7:**
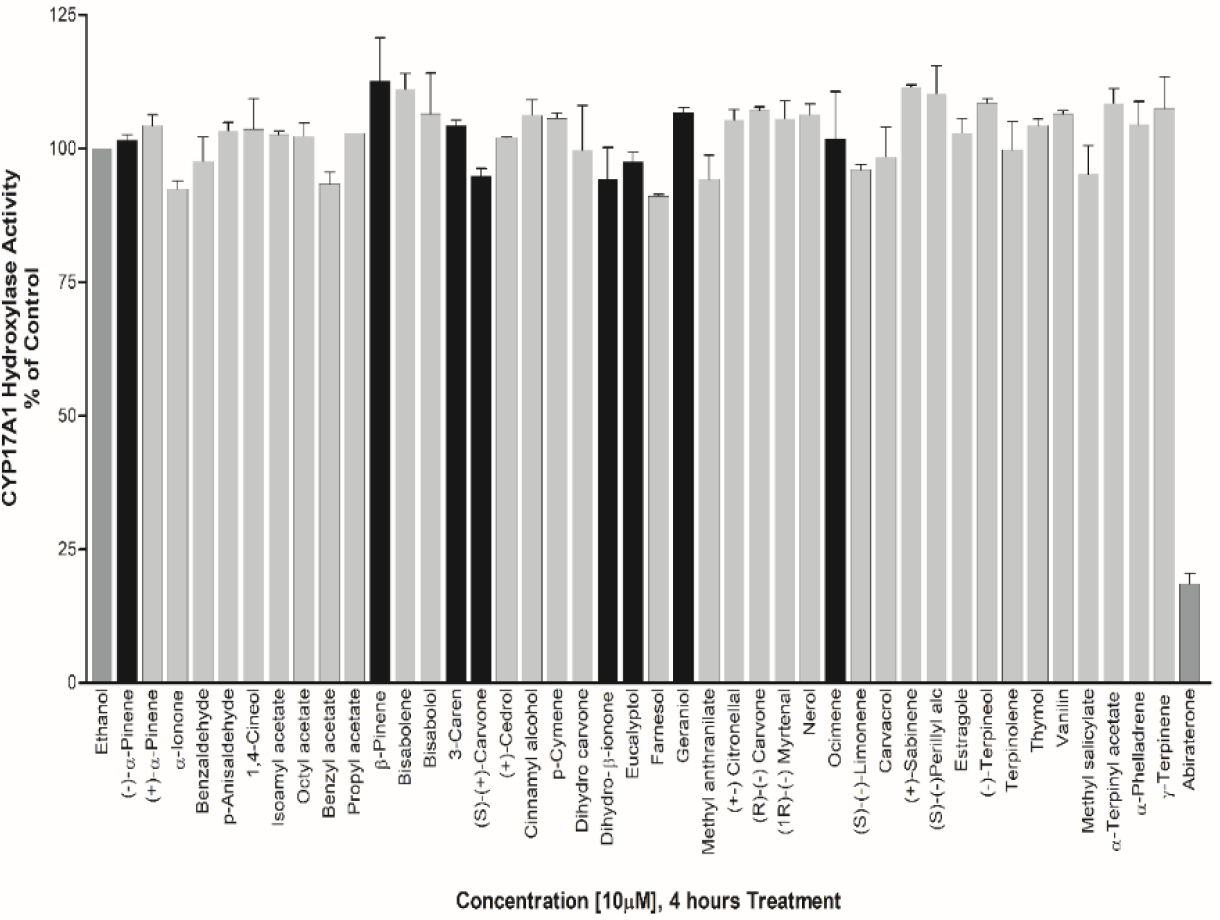
Assay of CYP17A1 17-hydroxylase activity. Essential oil compounds were tested for effects on CYP17A1 17-hydroxylase activity using radiolabelled progesterone as substrate and conversion to 17-hydroxy progesterone was monitored using autoradiography of steroids after separation by TLC. Effects were calculated as percentage of control.

However, EOs such as Eucalyptol, Geraniol, (S)-(+) Carvone, 3-Caren, Ocimene, β-Pinene, (-)-α-Pinene, Dihydro-β-ionone showed about 13%, 13%, 15.2%, 16%, 18%, 19%, 20% and 31% inhibition in CYP17A1 Lyase activity respectively. The effect of EOs towards an exclusive inhibition of CYP17A1 lyase activity makes them good candidates to study further as basic structural leads for designing more potent inhibitors (**Figure 8**).

**Figure 8:**
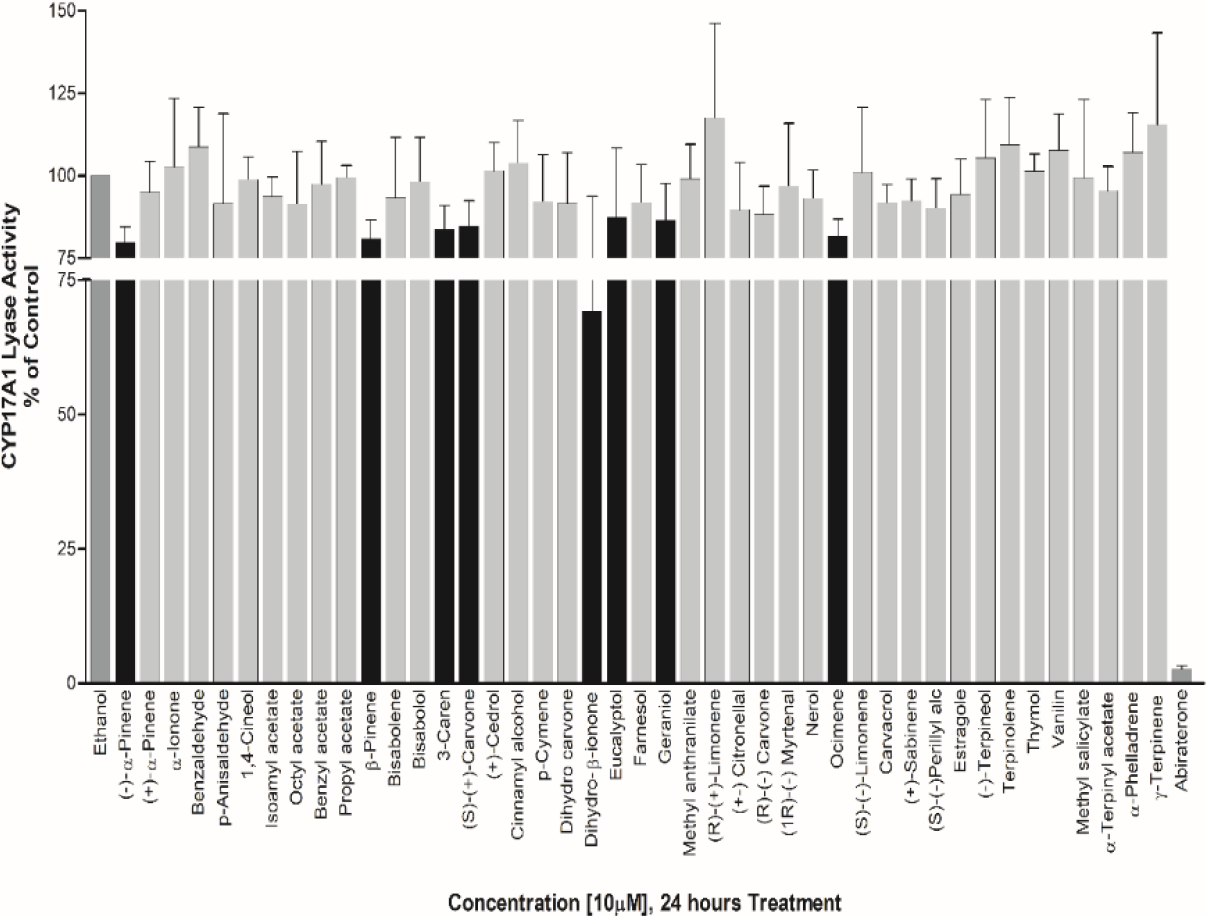
Effect of essential oil compounds of CYP17A1 17,20 lyase activity. Compounds were tested for effects on CYP17A1 17,20 lyase activity using radiolabeled 17OH-Pregnenolone as substrate and conversion to DHEA was monitored using scintillation counting. Effects were calculated as percentage of control.

### Effect on CYP19A1 Activity

EOs exhibiting significant effect on CYP17A1 activity and those predicted to be estrogenic in nature in some literatures were selected for screening of CYP19A1 activity. Bisabolol, Cedrol, Dihydro-β-ionone, (R)-(+)-Limonene, (-)-Terpineol, and α-Terpinyl acetate showed significant inhibition of aromatase at about 22%, 29%, 33%, 26%, 27%, and 29% respectively (**Figure 9**). Dihydro-β-ionone was found to be the most effective inhibitor of both CYP17A1 Lyase activity and CYP19A1 activity.

**Figure 9:**
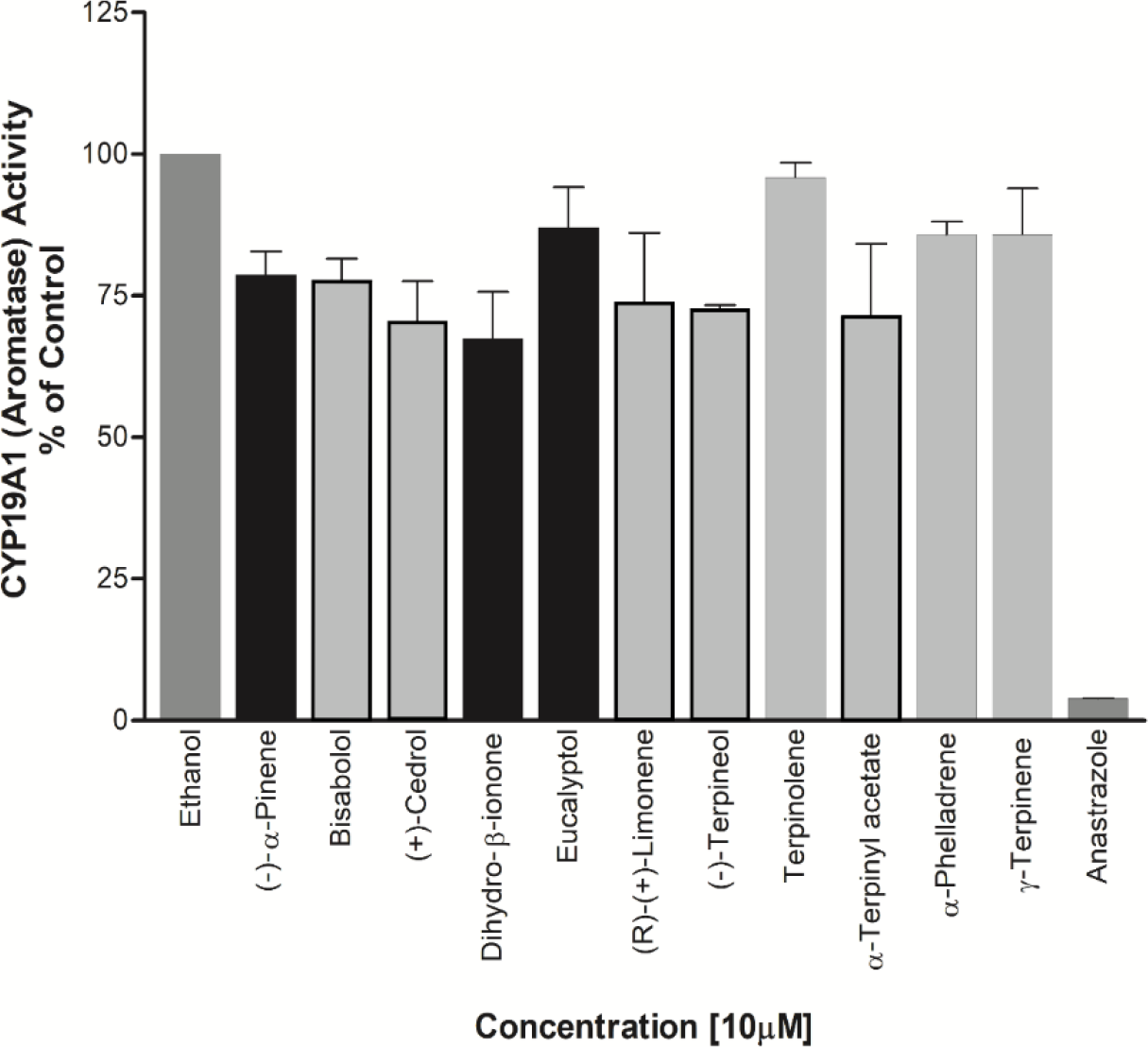
Effects of essential oil components on CYP19A1 activity. Essential oil components were tested against CYP19A1 activity using microsomes prepared from placental JEG-3 cells. Radiolabeled androstenedione was used as substrate and conversion to estrone was monitored by water release assay. A known CYP19A1 inhibitor anastrazole was used as positive control.

### Effect on Prostate Cancer Cell Viability

(-)-α-Pinene, Dihydro-β-ionone, Eucalyptol were also found to be causing cell toxicity in the prostate cancer cell line, LNCaP cells. All of these compounds showed an increased potency for cell growth inhibition with increasing treatment durations. Up to 50% reduction in cell viability was observed when the cells were treated with these EOs for 48 hours. Cedrol which showed maximum inhibition of CYP19A1 activity was also found to be reducing the cell viability of LNCaP cells. However, since a direct effect on CYP17A1 activity was not observed by cedrol, a different mechanism may be involved in toxicity towards LNCaP cells (**Figure 10**).

**Figure 10:**
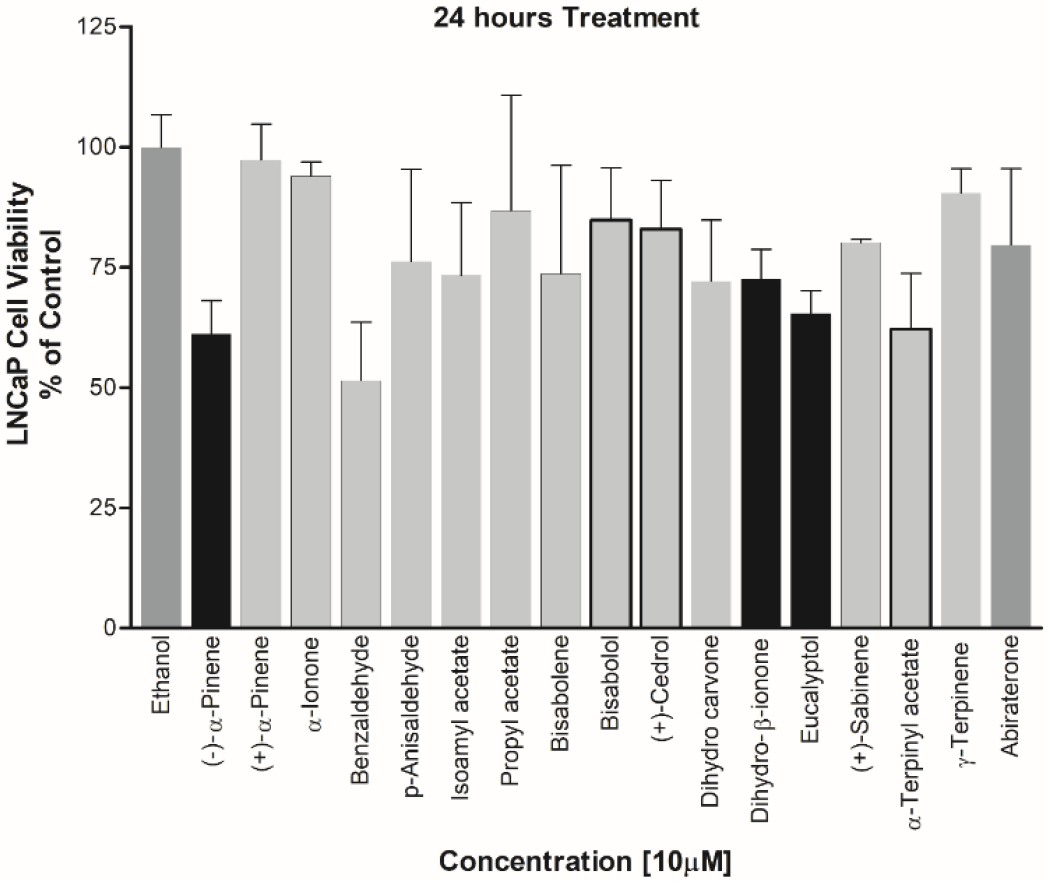
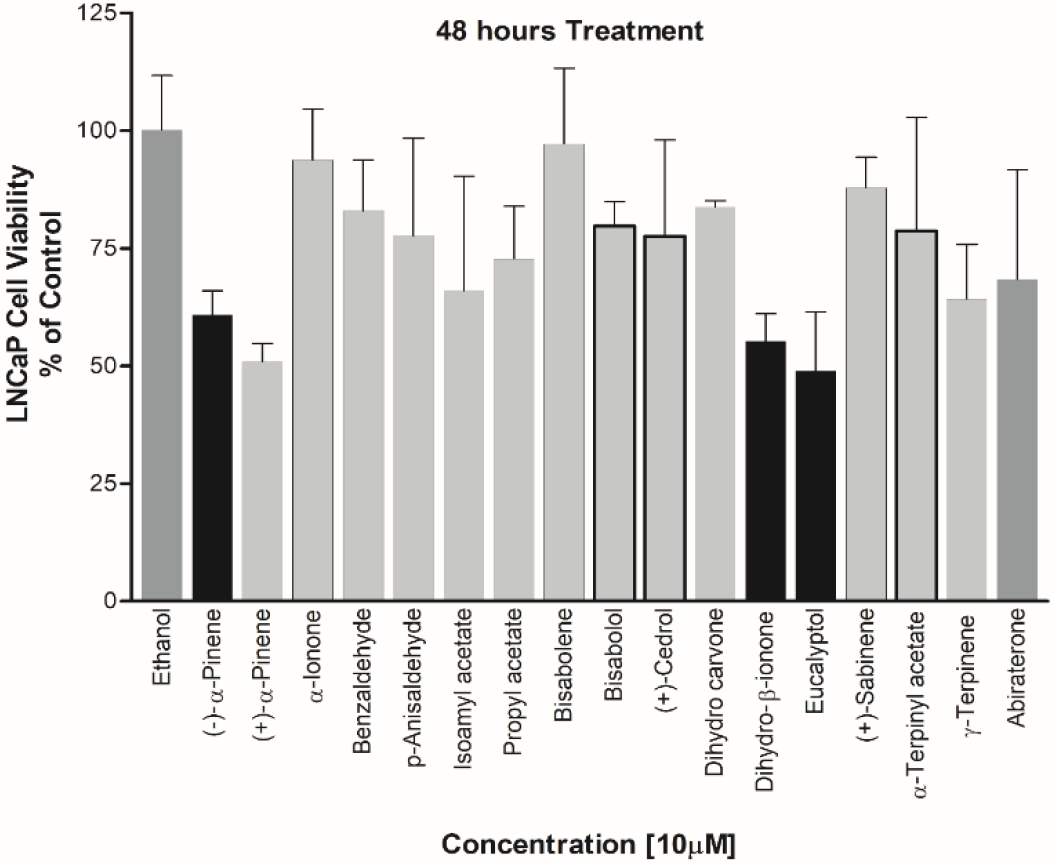
Effect of essential oil components on LnCaP cell proliferation. Essential oil compounds were checked for effect on proliferation of androgen dependent prostate cancer cell line, LnCaP cells. Cell viability was determined after 24h and 48h treatment with selected compounds that showed inhibitory effects in CYP17A1 or CYP19A1 assays.

### Steroid analysis by LC-MS/MS

Individual steroid levels were normalized to the amount of Pregnenolone (P5). P5 was the starting steroid substrate to profile all the steroids in the biosynthetic pathway. The addition of dihydro-β-ionone to adrenal cells did not alter the levels of 17OHP5 or DHEA, however DHEA-S levels appeared lower (about 8%) compared to the control. The addition of Eucalyptol reduced DHEA levels (about 13%), 1.2-fold (approaching significance at *p*=0.0574), also with lower DHEA-S levels compared to the control (**Figure 11**).

**Figure 11:**
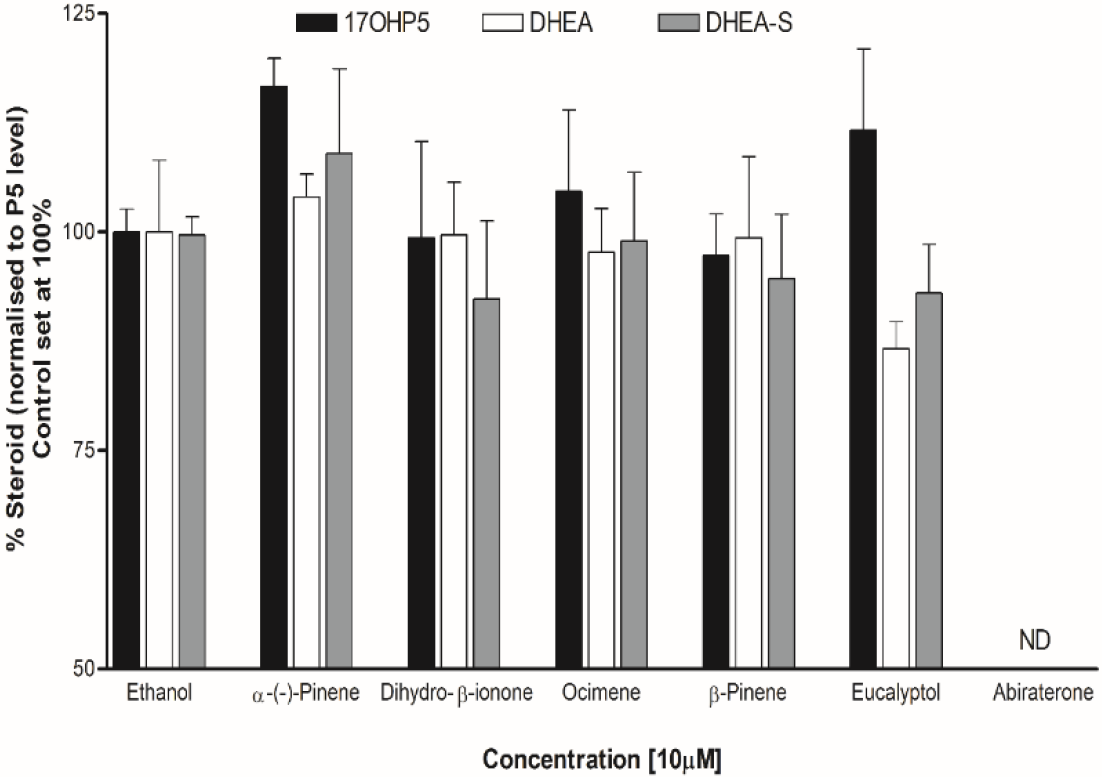
Steroid profiling using LC-MS/MS. Effect of essential oil components was tested for overall effects on steroid profiles of human adrenal NCI-H295R cells. Cells were treated with different compounds and steroid metabolites were analyzed by LC-MS/MS. Results shown are normalized against pregnenolone.

## 4. Discussion

Essential oils are highly concentrated plant extracts that are routinely used in in wellness, beauty, and cleaning products. However, since EOs are not pharmaceutical products, they are not regulated and therefore, their safety profiles are a topic of concern due to potential adverse reactions associated with their use. The safety of essential oils may also depend on individual metabolic profiles. In the past, few studies have been performed about the effect of EOs on cytochrome P450 enzymes that are involved in drug and xenobiotic metabolism [62]. Spicakova et. al. have studied the effect of sesquiterpenes beta-caryophyllene oxide and trans-nerolidol by docking into CYP3A4 and checking the results by functional assays of enzyme activity [63, 64]. Beta-caryophyllene oxide binds to CYP3A4 close to the heme without coordinating to the Fe atom and also showed a waek inhibition of CYP3A4 activity. Similarly a waek inhibition of CYP2C8, CYP2C9 and CYP2C19 was observed for cedrene, cedrol and thujopsene, but cedrol showed strong inhibition of CYP3A4 and CYP2B6 [65].

The steroidogenesis, leading to developmental and reproductive changes as well as impact on immunological and neurological changes linked to steroid hormones. Here we have investigated the impact of several essential oil components on steroid production mediated by CYP17A1 and CYP19A1, two key enzymes involved in regulation of androgen and estrogen in human. While essential oils offer potential benefits, their use should be approached with caution due to potential safety issues and hormonal imbalances. It is crucial to conduct thorough research and consult healthcare professionals before incorporating essential oils into daily routines.

On the other hand, anti-androgen properties of essential oil compounds may offer structural leads for the design of novel drugs targeting steroid hormone production in metabolic disorders dependent on steroid hormones, including prostate cancer, breast cancer and poly cystic ovary syndrome.

Currently, the classical methods employed in the treatment of PCa include Androgen Deprivation Therapy using CYP17A1 inhibitors and/or blocking AR binding to its ligand using AR antagonists [66] [67] (**Figure 12**). CYP17A1 targeted drugs have been developed over the years for the treatment of PCa as well as Castration Resistance Prostate Cancer (CRPC) [68]. Therefore, CYP17A1 has emerged as an attractive target for the design of inhibitors to use as drugs against prostate cancer [69]. From a non-selective Cytochrome P450 inhibitor, Ketoconazole, first-generation CYP17A1 targeted drugs such as Abiraterone and Orteronel (TAK700) to the most recent compounds with better selectivity towards 17,20 Lyase activity like Galeterone (TOK-001) and VT464, there is a continuous search for more efficient and potent inhibitors to overcome the challenges due to their adverse side-effects [70] [71] [72] [73] [74] [75-78]. For instance, in addition to CYP17A1, Abiraterone also targets cytochrome P450 21-hydroxylase (CYP21A2) activity, which is essential for aldosterone and cortisol production [79, 80]. As a result, suboptimal levels of cortisol due to inhibition of CYP21A1 leads to the requirement of glucocorticoid co-therapy in these patients [81]. However, most treatments have only a small effect in improving PCa patient survival. Next-generation drugs like Galeterone acts as both an AR antagonist and a CYP17A1 inhibitor. However, in metastasized PCa conditions such as CRPC, the tumor cells develop mechanisms to evade androgen dependency for their growth and survival. These mechanisms include the expression of AR variants which bind to androgen precursors of adrenal origin and *de novo* intra-tumoral androgen production [82] [83]. Current strategies for the development of drugs focus on designing inhibitors with the ability to modulate the elevated levels of circulating androgens as well as steroids derived from alternative pathways in the case of androgen independent PCa without disturbing the cortisol metabolism [84] [85]. Therefore, in the search for novel inhibitors of CYP17A1 with improved target specificity and reduced off-target effects, EOs could be utilized as chemical leads for the designing novel drugs against PCa and PCOS.

**Figure 12:**
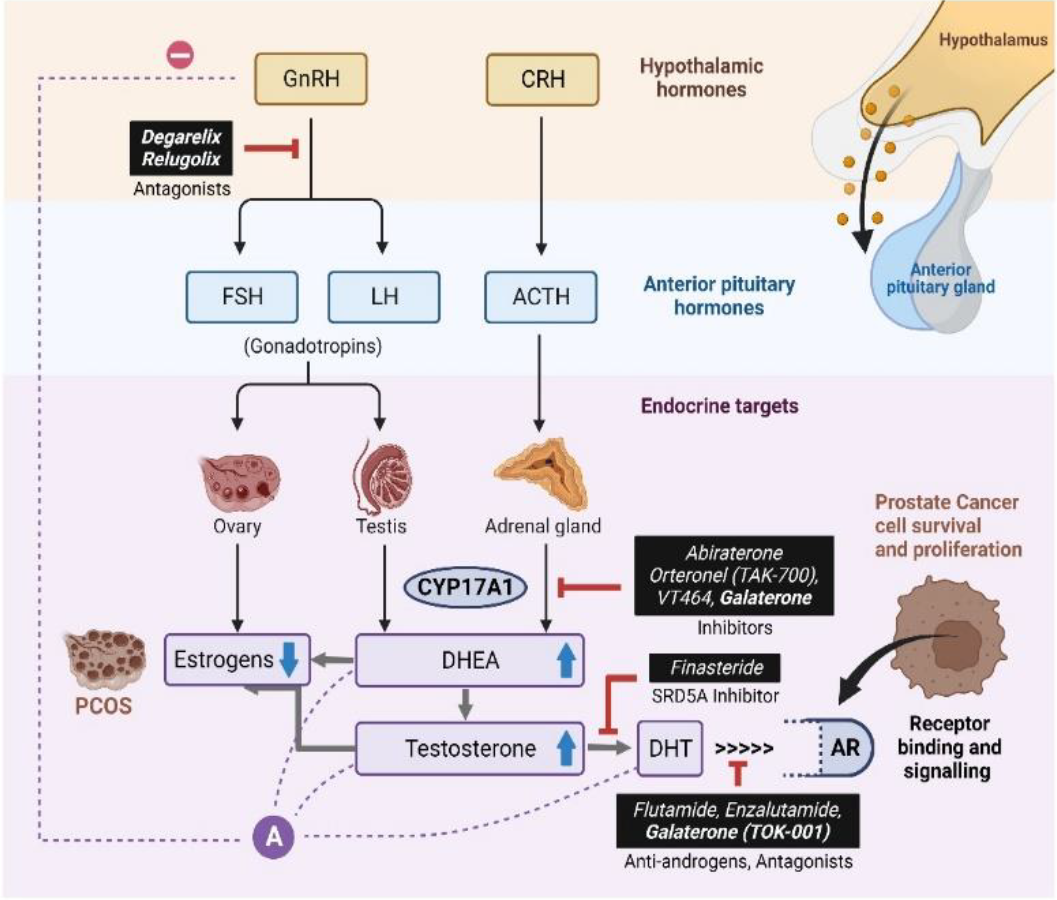
Different approaches to target androgen production for prostate cancer treatment.

## Supplementary Materials

The following supporting information can be downloaded at: www.mdpi.com/xxx/s1, File S1:. List of compounds used in this study.

## Author Contributions

Conceptualization, A.V.P., A.L.; K.S.; CYP17A1 assays, K.S.; cell viability, K.S., J.Y., and S.T.; steroid profiling, T.d.T. and C.D.V.; molecular modeling, S.T. and F.S.J.; writing—original draft preparation, KS and AVP.; writing—review and editing, T.d.T., C.D.V., F.S.J., A.V.P., project administration AVP. All authors have read and agreed to the published version of the manuscript.

